# Single-agent PD-L1 blockade and tumour volume in orthotopic GL261 glioma, with an exploratory lipid analysis of the treated tumours

**DOI:** 10.64898/2026.11.22.768425

**Authors:** Paula Carretero-Navarro, Pilar López-Larrubia, Jesús Pacheco-Torres

## Abstract

Immune checkpoint blockade has not yet changed outcomes in glioblastoma (GBM), and in immunocompetent mouse glioma the effect of a single blocking antibody is usually reported as survival rather than as tumour volume with its own confidence interval. We treated mice bearing orthotopic GL261 gliomas with anti-PD-L1 as a single agent in an IDH1-wild-type and in an IDH1-mutant line, and followed tumour volume by MRI. At day 14, tumours of treated animals were 0.33 times the volume of control tumours in the IDH1-wild-type line (95 % CI 0.21 to 0.53; p < 0.001; 16 anti-PD-L1 and 20 control animals) and 0.42 times in the IDH1-mutant line (0.22 to 0.78; p = 0.006; 10 and 10 animals), from a linear mixed model of log tumour volume over days 8 to 16. Diffusion imaging of the same tumours gave a higher apparent diffusion coefficient in the treated IDH1-wild-type tumours, by 23 % over the whole tumour (p = 0.027), and lower fractional anisotropy in the treated IDH1-mutant tumours, by 25 % over the whole tumour (p = 0.023), and in the periphery of the IDH1-wild-type ones, by 24 % (p = 0.005). We also imaged the lipid composition of treated and control IDH1-wild-type tumours ex vivo by mass spectrometry imaging: relative to the healthy brain of the same section, sphingomyelin was higher in the treated tumours and phosphatidylcholine species shifted towards shorter chains (p = 0.016 for both). In orthotopic GL261 glioma, anti-PD-L1 as a single agent reduces tumour volume in the IDH1-wild-type and in the IDH1-mutant line.

## Introduction

Glioblastoma is the most common malignant primary brain tumour in adults, and the standard of care established two decades ago, radiotherapy with concomitant and adjuvant temozolomide, raised median survival from 12.1 to 14.6 months [1]. Immune checkpoint blockade has changed outcomes in several solid tumours, and it has so far not done it in glioblastoma. In the only phase 3 trial of a PD-1 inhibitor in recurrent glioblastoma, nivolumab did not improve overall survival over bevacizumab, with medians of 9.8 and 10.0 months [2]. The schedule, however, appears to matter, as neoadjuvant pembrolizumab extended survival over the same drug given only after surgery in a randomised trial of 35 patients [3]. What in the tumour, the host or the timing accounts for that difference is not settled.

PD-L1 is detectable in the large majority of glioblastoma specimens, alongside tumour-infiltrating lymphocytes, and published series disagree on whether the level of that expression carries prognostic information, one finding no relation to outcome [4] and another finding higher expression associated with shorter survival [5]. In immunocompetent mouse models the preclinical picture is also less uniform than it is usually presented. In orthotopic GL261 glioma, anti-PD-1 as a single agent did not prolong survival, and long-term survival was reached only when the antibody was combined with stereotactic radiation [6]. In the same model, a systematic comparison of blocking antibodies reported long-term tumour-free survival in 50 % of animals treated with anti-PD-1 and in 20 % of those treated with anti-PD-L1 [7]. GL261 is moderately immunogenic [8] and carries a high mutational load, and it responds to checkpoint blockade where a model with a low mutational load does not [9]. What a single antibody does to tumour volume in this model has seldom been reported.

In the 2021 World Health Organization classification, the adult-type diffuse gliomas comprise only three types, and the name glioblastoma now belongs to the IDH-wildtype one alone, the designation glioblastoma, IDH-mutant having been eliminated [10]. The IDH1 R132H mutation, in addition to metabolic changes, also shapes the immune compartment of the tumour that carries it. Mutant IDH1 lowers STAT1 and the interferon-inducible chemokine CXCL10 and reduces the accumulation of CD8-positive T cells at the tumour site [11]. The oncometabolite R-2-hydroxyglutarate that mutant IDH produces is taken up by T cells and suppresses their activity, and IDH1-mutant gliomas contain fewer T cells [12]. The same metabolite inhibits complement activation and the migration and proliferation of activated T cells [13]. How a glioma line carrying this mutation behaves under single-agent PD-L1 blockade has been examined far less often than the wild-type case.

Tumour lipid metabolism offers a second and independent window on the same tumours. Abnormal choline metabolism is a feature of malignant transformation, in which the precursors and breakdown products of membrane phospholipids rise and become detectable by magnetic resonance spectroscopy [14]. Matrix-assisted laser desorption ionisation mass spectrometry imaging measures lipid classes in a tissue section without losing their position, and in orthotopic mouse glioma it has resolved changes in phosphatidylcholine and sphingolipid species between tumour and surrounding brain [15]. Whether single-agent PD-L1 blockade leaves any trace in that lipid composition is the question that the exploratory part of this work sets out to address. Here we report tumour volumes measured by MRI in mice bearing orthotopic GL261 gliomas, in the IDH1-wild-type and in the IDH1-mutant line. Animals received anti-PD-L1 as a single agent, and the two lines were analysed separately throughout. We also report an exploratory analysis of lipid classes by mass spectrometry imaging in a subset of the wild-type animals.

## Methods

### Animals and ethics

All procedures were authorised under reference PROEX 292.0/22 by the Dirección General de Agricultura, Ganadería y Alimentación of the Comunidad de Madrid, on a report of its Animal Protection Area, for the user centre Instituto de Investigaciones Biomédicas Alberto Sols (ES280790000188). They were carried out in accordance with Spanish Royal Decree 53/2013, which transposes Directive 2010/63/EU on the protection of animals used for scientific purposes. Male and female C57BL/6j mice, aged 10 to 12 weeks at implantation, were bred in the animal facility of the same centre. Animals were housed in groups, in acoustically isolated rooms under a controlled light and dark cycle, with stable temperature and humidity, environmental enrichment and daily observation. Both sexes were included in every arm of both genotypes, and the sex of each animal was taken from the laboratory record. Of the animals analysed here, 18 were female and 18 were male in the IDH1-wild-type line, and 10 were female and 10 were male in the IDH1-mutant line.

### Tumour model

Tumours were established by orthotopic implantation of GL261 cells, either wild-type or stably expressing the IDH1 R132H mutation, into the brain of recipient mice. Anaesthesia was induced and maintained with 1 to 2 % isoflurane in oxygen, and the animal was held in a stereotactic frame. The scalp was shaved and disinfected with 70 % ethanol and povidone iodine, a midline incision was made, and a parietal burr hole was drilled with a 25G needle 2.3 mm to the right of Bregma. Either 10,000 wild-type cells or 100,000 IDH1 R132H cells, in 4 µL of culture medium, were injected at a depth of 3.3 mm through a 27G needle on a glass Hamilton syringe mounted on the frame. The burr hole was sealed with surgical cement and the skin was closed with 3/0 braided silk. Meloxicam at 2 mg/kg and buprenorphine at 0.1 mg/kg were given subcutaneously before surgery and on the three following days. The two lines were implanted, imaged and analysed separately, and they were never pooled or compared with each other.

### Treatment

Anti-PD-L1 antibody, clone 10F.9G2 (rat IgG2b κ; Bio X Cell BE0101, RRID AB_10949073), was given as a single agent by intraperitoneal injection. The first dose of 500 µg was given on day 6 or 7 after implantation, and 250 µg were given every three days thereafter, until death or the humane endpoint. The control group comprised animals receiving an isotype control antibody and animals receiving saline. The isotype control was rat IgG2b anti-KLH, clone LTF-2 (Bio X Cell BE0090, RRID AB_1107780), given at the same dose and on the same schedule as the anti-PD-L1 antibody. Saline was given in the same volume and by the same route.

### Which animals entered the analysis

An animal entered the growth analysis if it had at least one scan between days 8 and 16 after implantation, and every scan of that animal inside that window was then included. This gave 16 anti-PD-L1 and 20 control animals in the IDH1-wild-type line, over 89 scans, and 10 anti-PD-L1 and 10 control animals in the IDH1-mutant line, over 26 scans. Animals given saline and animals given an isotype control antibody were pooled into one control group; their tumours did not differ in growth in either line (p ≥ 0.27).

### MRI acquisition and volumetry

Tumours were followed in vivo on a Bruker BioSpec 70/16 operating at 7 T under ParaVision 6.0.1. Anaesthesia was induced with 4 % isoflurane in oxygen and maintained at 1 to 2 % through a nose cone, with a recirculating warm water blanket, a respiratory sensor and a rectal probe, the respiratory rate guiding the depth of anaesthesia. Volumetry was performed on a T2-weighted RARE series with a RARE factor of 8, a repetition time of 2,500 ms and an echo time of 8.70 ms, giving an effective echo time of 26.1 ms. The field of view was 18 × 18 mm on a 200 × 200 matrix, that is 0.09 × 0.09 mm in plane, with 12 slices of 0.8 mm and 3 averages. The tumour was outlined slice by slice on those images, manually and independently by two people, and the volume was obtained from the outlined area and the slice geometry. The images shown in Figure 1A are representative sections chosen for similar tumour volume.

**Figure 1.**
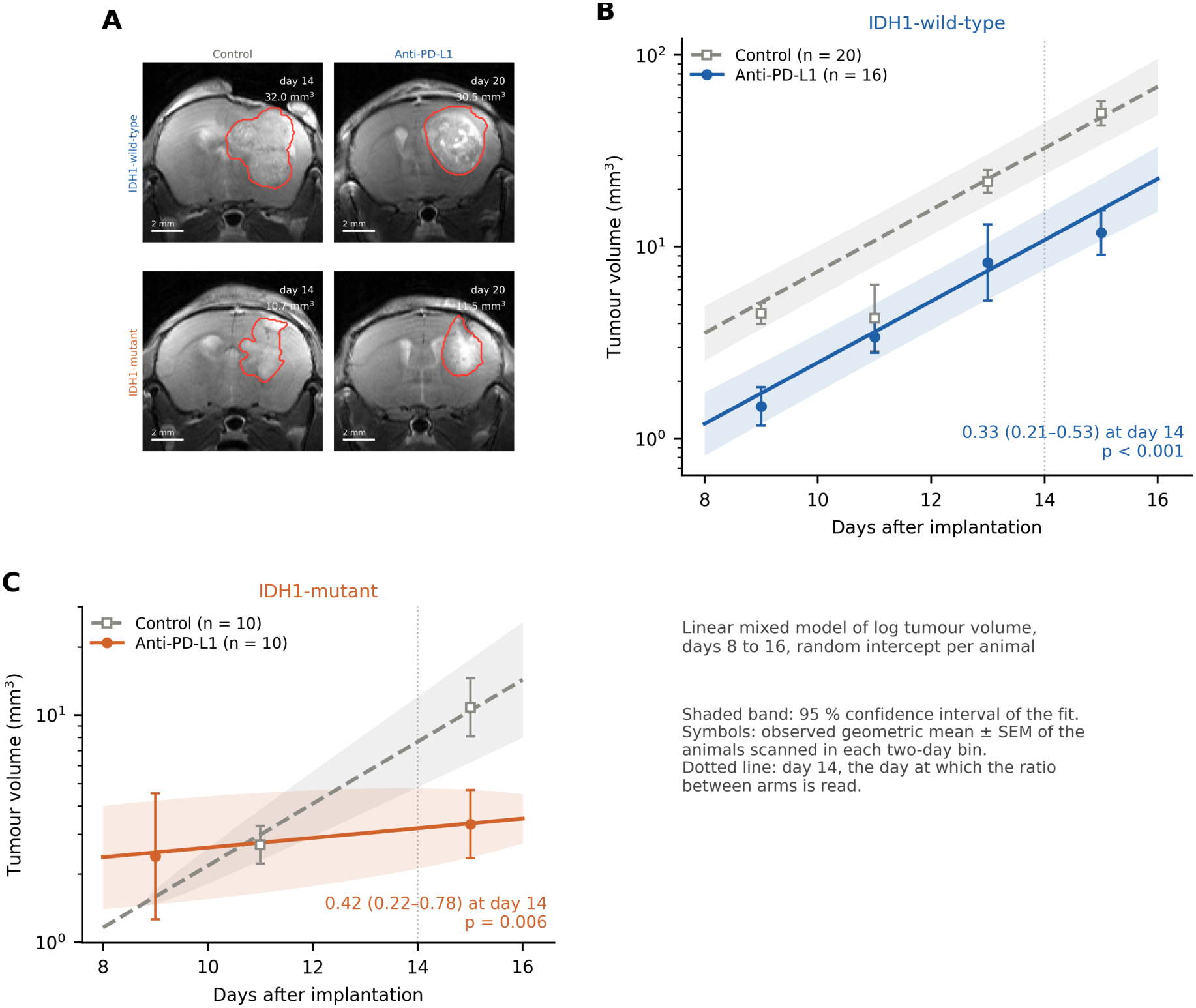
Tumour volume under single-agent anti-PD-L1 in IDH1-wild-type and IDH1-mutant GL261 glioma. **(A)** T2-weighted sections with the outline used for volumetry drawn on the tumour, for a control animal and an anti-PD-L1 animal of similar tumour volume in each line: IDH1-wild-type above, IDH1-mutant below. The sections are matched by tumour volume and not by day, and in both lines the tumour of the treated animal reaches an equivalent volume six days later than that of the control. Each image carries its day and its volume. Scale bars, 2 mm. **(B)** Tumour volume against day after implantation in the IDH1-wild-type line, on a logarithmic scale. The lines are the fit of the mixed model of log tumour volume over days 8 to 16 and the bands its 95 % confidence interval, grey for control and blue for anti-PD-L1; the symbols are the observed geometric mean and its standard error of the animals scanned in each two-day bin. The dotted line marks day 14, at which the ratio between arms is read. **(C)** The same for the IDH1-mutant line, in orange. n = 16 anti-PD-L1 and 20 control animals in the IDH1-wild-type line, over 89 scans, and 10 and 10 animals in the IDH1-mutant line, over 26 scans. The ratio between arms at day 14, its 95 % confidence interval and its p value are printed in each panel.

### Diffusion MRI

Diffusion-weighted images were acquired in the same sessions, in 30 directions at b values of 800 and 2,500 s/mm^2^ together with 3 images without diffusion weighting. The gradient duration was 4 ms and the gradient separation 25 ms, with a repetition time of 3,000 ms, an echo time of 38.1 ms and 3 averages. The field of view was 18 × 18 mm on a 100 × 100 matrix, with 12 slices of 0.8 mm. The apparent diffusion coefficient and the fractional anisotropy were taken from the tensor fit of one imaging session per animal. Each tumour was divided on the T2-weighted images into a core and a peripheral region, and the same contours were applied to the diffusion maps, so that each animal contributes one value per measure and region. A quality criterion set in advance and applied identically to every scan removed measurements whose apparent diffusion coefficient or mean diffusivity fell outside the median ± 3.5 median absolute deviations of the series, and regions of fewer than 100 voxels were not used. This left 7 anti-PD-L1 and 8 control animals in the IDH1-wild-type line and 5 and 5 in the IDH1-mutant line.

### Statistical analysis

The unit of analysis is the animal, not the scan and not the voxel. Tumour growth was modelled with a linear mixed model of the natural logarithm of tumour volume against day after implantation and treatment, over days 8 to 16, with a random intercept per animal; in the IDH1-wild-type line a fixed term for the series of implantations was added. Time was centred on day 14, so that the treatment coefficient is the difference between arms at that day and the ratio we report is its exponential, with Wald 95 % confidence intervals. Day 14 is a reference point of the model and not a day on which every animal was measured. One volume measurement that was physiologically impossible was excluded before fitting, and the animal that provided it stays in the analysis with its remaining scans. In the IDH1-mutant line most animals contribute a single scan inside that window and the fit returns a convergence warning. The estimate is unchanged by other optimisers and by a regression with errors clustered by animal, which give a ratio of 0.41 with p between 0.005 and 0.010.

The effect of sex was examined in the IDH1-wild-type line by fitting the same model in females and in males and by adding sex and its interaction with treatment. In the IDH1-mutant line the sex of the animals coincides with the order in which they were implanted, so the two cannot be separated and no analysis by sex is reported. The two lines were analysed separately throughout, and they were neither pooled nor compared with each other.

Each diffusion measure was compared between arms region by region with Welch’s t-test, in each line separately, and the effect is given as a percentage of the control mean. The diffusion analysis was exploratory.

No correction for multiple comparisons was applied to the growth or the diffusion analyses, and the p values reported for them are uncorrected. Analyses were run in Python 3.8.5 with statsmodels 0.14.1, pandas 2.0.3, numpy 1.23.5 and scipy 1.10.1.

### MALDI mass spectrometry imaging

This section describes the exploratory lipid imaging performed on four anti-PD-L1 animals and five control animals of the wild-type line, one cryosection per animal. The four anti-PD-L1 animals are four of the treated wild-type animals of the growth analysis, and their brains come from their final imaging session, on days 19 to 20 after implantation.

Animals were perfused with 1× PBS before the brain was removed. Brains were frozen immediately in liquid nitrogen and kept at −80 °C until sectioning. Sections of 12 µm were cut on a Leica CM1950 Dual EcoTec cryostat with chamber and blade head at −20 °C, using a small base of OCT placed on the cerebellum so that it never touched the tissue of interest, and were thaw-mounted onto MALDI IntelliSlides (Bruker). Mounted sections were stored at −80 °C until analysis and were dried in a vacuum desiccator in the dark for 45 min immediately before matrix application.

Norharmane matrix was prepared fresh at 7 mg/mL in chloroform:methanol 2:1 (v/v) and sonicated for 5 min. It was deposited with an HTX TM-Sprayer at 30 °C, a flow rate of 0.05 mL/min, a nozzle velocity of 1,200 mm/min and 14 passes. Coated sections were dried again in a vacuum desiccator in the dark for 15 min before acquisition.

Images were acquired on a timsTOF fleX (Bruker) under timsTOF 6.0.6 with the method lipids_tims_on_imaging_50um_pos.m, in positive ion mode over m/z 300 to 1,350, at a pixel size of 50 µm with a 46 × 46 µm raster. The laser was operated at 10 kHz with 200 shots per pixel and 25 % power. Trapped ion mobility was active over 1/K0 from 0.7 to 1.8 V·s·cm^−2^, with a 200 ms ramp and 20 ms accumulation. The instrument was calibrated with ESI-L Low Concentration Tuning Mix (Agilent). Sections of treated and of control animals were acquired in separate sessions, with the same instrument and the same method and each against its own calibration. Images were exported from SCiLS Lab 14.01 as imzML and read with software written for this work in Python.

The same sections then entered a sequential workflow, in which the matrix was removed and tryptic digestion was performed for peptide imaging. The lipid images reported here were acquired first, before any washing step, so that nothing done afterwards can affect them. No peptide data are used in this work.

Ion intensities were integrated within ±10 ppm of the m/z of each feature, without recalibration. For annotation only, m/z values were corrected for a −3.3 ppm offset estimated from 108 unambiguous phosphatidylcholine [M+H]+ ions, and were matched to sum-composition lipid species as [M+H]+, [M+Na]+ and [M+K]+, with ion mobility used where available. Species are therefore assignments by accurate mass and ion mobility, without tandem mass spectrometry confirmation. Free phosphocholine (m/z 184.07) lies below the acquired mass range and was not measured.

Each pixel was normalised to its root mean square before class intensities were summed, which is the normalisation recommended for heterogeneous tissue because it is robust to dominant peaks. Every value reported for a lipid class is therefore a relative abundance, a fraction of the normalised spectrum, and every comparison is expressed as a log2 ratio against a reference region of the same section. That reference is the healthy brain of the same section, defined as all tissue more than 300 µm from the tumour in both hemispheres, and it is the same reference in every panel of Figure 3.

Two measures of sphingomyelin in the tumour are reported. The first is the median of the per-pixel log2 index over the tumour. The second is the fraction of tumour pixels whose index exceeds 0.5 in log2 units, that is 1.4 times the healthy brain, a threshold that was chosen when the figure was drawn and not before.

The unit of analysis is the animal, not the pixel, and each animal contributes one value per measure. Treated and control animals were compared by an exact permutation test of the group labels, with the difference of means as the statistic, over the 126 ways of choosing 4 animals from the 9, two-sided. Because the treated tumours were smaller, each measure was also adjusted for tumour size in a linear regression of the value on treatment and on the log2 of the tumour area in pixels. That regression was fitted across the 13 wild-type animals imaged, which include four animals of the AGI-5198 and 1-methyl-tryptophan arms of the same series. Those four contribute only to the estimate of the relationship with size, and they are not compared with the treated animals. Its p value is an exact permutation of the anti-PD-L1 label among the 13 animals, over 715 splits, two-sided.

The smallest p value attainable with four treated and five control animals is 0.008, and p values are reported uncorrected because the analysis is exploratory. Across the 17 class-level tests, Benjamini-Hochberg correction leaves no class below 5 %, the lowest q value being 0.07. Treated and control sections were acquired in different sessions, on the same instrument and with the same method, which is why every value is a ratio against a reference region of the same section rather than an absolute intensity. The result is unchanged when the single control section acquired in the same session as the treated sections is left out.

## Results

### Tumour growth

To test whether single-agent anti-PD-L1 reduces tumour volume in orthotopic GL261 glioma, we followed IDH1-wild-type tumours by MRI (Figure 1A) and modelled log tumour volume over days 8 to 16 (Figure 1B). At day 14, tumours of treated animals were 0.33 times the volume of control tumours (95 % CI 0.21 to 0.53; p < 0.001; 16 anti-PD-L1 and 20 control animals, 89 scans). The antibody given alone was therefore sufficient to lower tumour volume in this line.

To examine whether the response depended on sex, the same model was fitted in females and in males. The ratio at day 14 was 0.41 in females (95 % CI 0.26 to 0.65) and 0.25 in males (0.12 to 0.52), and the interaction between sex and treatment gave 0.89 (0.37 to 2.10; p = 0.78), so the response did not differ between sexes in this line.

The same measurements and the same model were applied to tumours of the IDH1-mutant line (Figure 1C). At day 14, tumours of treated animals were 0.42 times the volume of control tumours (95 % CI 0.22 to 0.78; p = 0.006; 10 anti-PD-L1 and 10 control animals, 26 scans). In this line the sex of the animals cannot be separated from the order in which they were implanted, and no analysis by sex is reported.

### Water diffusion inside the tumour

To ask whether treatment also changed the tissue inside the tumour, each tumour was divided into a core and a peripheral region on the T2-weighted images, the same contours were applied to the diffusion maps, and the arms were compared region by region (Figure 2A). In IDH1-wild-type tumours the apparent diffusion coefficient was higher in treated animals, by 23 % over the whole tumour (p = 0.027) and by 29 % in the core (p = 0.030), and by 8 % in the periphery, which this comparison does not separate from zero (p = 0.17; 7 anti-PD-L1 and 8 control animals). Fractional anisotropy was 24 % lower in the periphery of those same tumours (p = 0.005), and 17 % lower over the whole tumour (p = 0.051).

**Figure 2.**
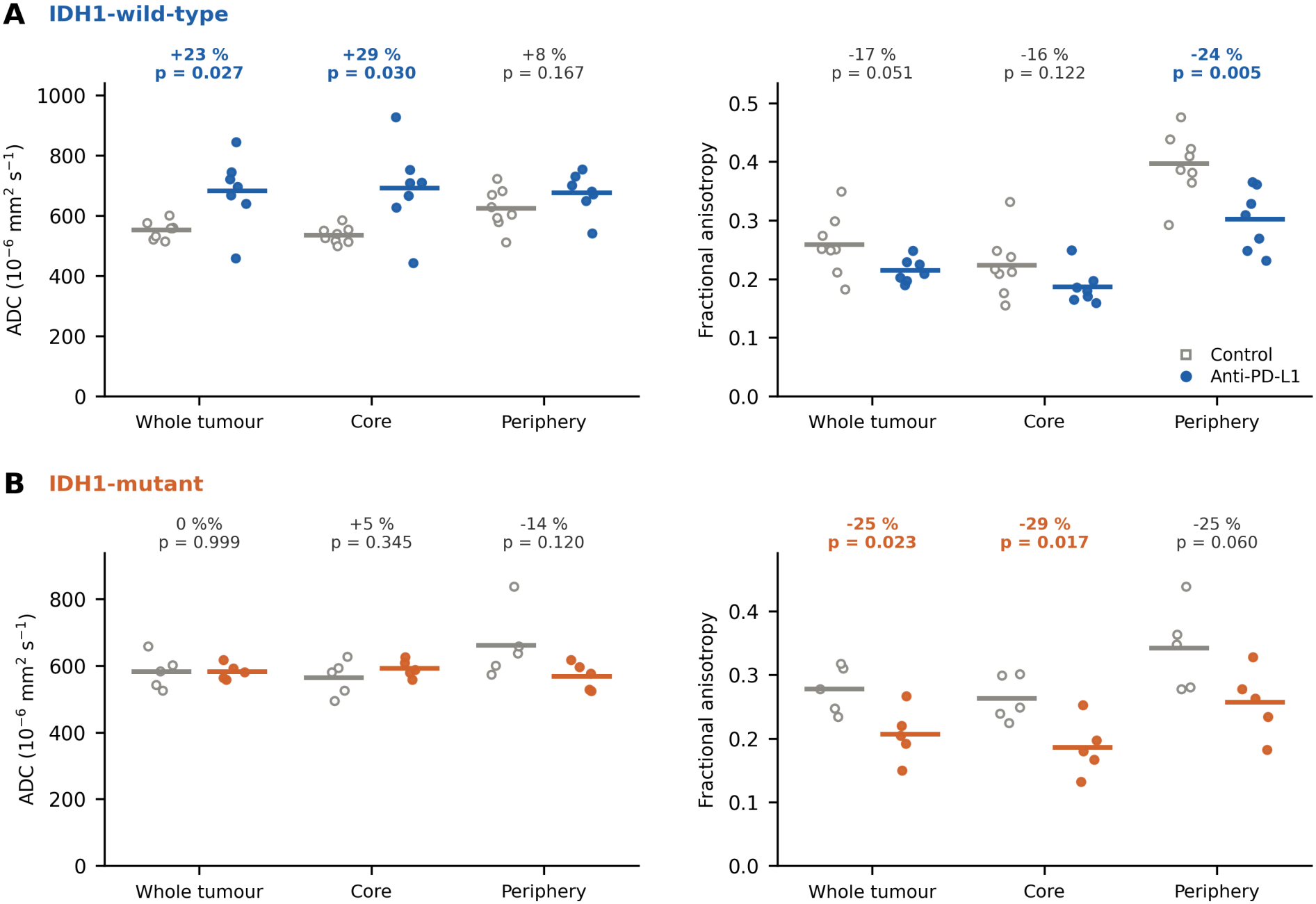
Water diffusion inside the tumour, by region. **(A)** IDH1-wild-type line. **(B)** IDH1-mutant line. In each panel, the apparent diffusion coefficient on the left and the fractional anisotropy on the right, each given for the whole tumour, for the core and for the periphery. One point per animal, open symbols for control animals and filled for anti-PD-L1, with the group mean drawn as a bar. Above each pair is the difference between the arms as a percentage of the control mean, with its p value from Welch’s t-test; the differences with p < 0.05 are in bold. The apparent diffusion coefficient is in 10^−6^ mm^2^ s^−1^ and the fractional anisotropy is dimensionless. n = 7 anti-PD-L1 and 8 control animals in the IDH1-wild-type line, and 5 and 5 in the IDH1-mutant line. One value per animal and region.

In the IDH1-mutant line the change was in fractional anisotropy rather than in the apparent diffusion coefficient (Figure 2B). Fractional anisotropy was 25 % lower in treated animals over the whole tumour (p = 0.023) and 29 % lower in the core (p = 0.017), while the apparent diffusion coefficient differed in no region (p ≥ 0.12; 5 anti-PD-L1 and 5 control animals).

### Lipid composition of the IDH1-wild-type tumours

To explore whether treatment left a trace in the lipid composition of the tumour, we imaged one cryosection from each of four anti-PD-L1 and five control wild-type animals by MALDI mass spectrometry imaging (MSI). Every lipid class was expressed as a ratio to the healthy brain of the same section. Sphingomyelin was the class that separated the two groups, and it did so by three measures of the same sections. Across the whole tumour its summed relative abundance was higher in treated than in control animals by 0.54 in log2 units, a rise of 45 % (p = 0.016, and p = 0.008 after adjustment for tumour size; Figure 3A, C). The median of the per-pixel index, which uses no threshold, was higher by 0.59 in log2 units (p = 0.016, and p = 0.004 after adjustment for size). The rise also occupied area. Pixels high in sphingomyelin covered 73 to 92 % of the tumour in three of the four treated animals and 7 to 18 % in the five controls, the fourth treated animal reaching 17 % (Figure 3B). That percentage differed between the groups with p = 0.024, and with p = 0.008 after adjustment for size. The rise was spread across the class rather than carried by one species, with 14 of the 16 cleanly resolved sphingomyelin species higher in treated animals. It did not follow markers of myelin (rho −0.09 and 0.17), which argues against admixed white matter.

**Figure 3.**
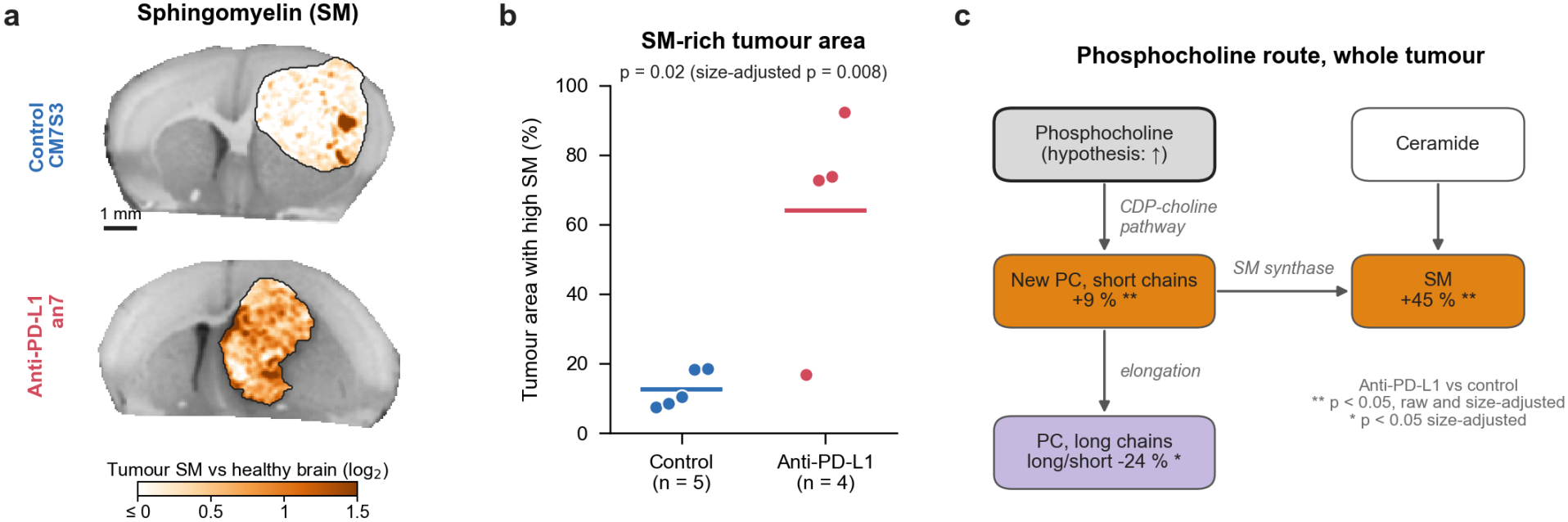
Sphingomyelin and phosphatidylcholine of IDH1-wild-type GL261 tumours under single-agent anti-PD-L1, by MALDI mass spectrometry imaging. One cryosection from each of nine animals, four treated with anti-PD-L1 and five controls. n are animals, never pixels. **(A)** Maps of species assigned to sphingomyelin in one control and one treated tumour of similar size. The tumour is outlined and shown in colour as the log2 ratio to non-tumoural brain of the same section, and the rest of the section is in grey. The colour scale starts at zero, so white marks tumour at or below the level of non-tumoural brain. Scale bar, 1 mm. **(B)** Percentage of the tumour area high in sphingomyelin, defined as more than 1.4 times the non-tumoural brain of the same section, with one point per animal and the group mean as a horizontal line. The p value printed in the panel belongs to this percentage, and it is 0.024 before and 0.008 after adjustment for tumour size. **(C)** Reactions that connect the lipid classes measured in the whole tumour, each box carrying the difference between treated and control animals as a fold change

Phosphatidylcholine moved in the other direction and towards shorter chains. The short-chain [M+H]+ series rose by 0.13 and the long-chain [M+K]+ series fell by 0.57 in log2 units. The long-to-short ratios of the two series fell by 0.40 and 0.45 (p = 0.056 and p = 0.032), and by 0.28 and 0.35 after adjustment for tumour size (p = 0.014 and p = 0.018; Figure 3C). Because each class is measured as a fraction of the normalised spectrum, the simultaneous falls in other classes are not independent decreases, and these ratios between classes are the measures that do not depend on that normalisation. The change was also present at the tumour border. There sphingomyelin was higher in treated than in control animals by 0.37 in log2 units against the peritumoral reference of the same section. All four treated animals lay above all five controls, with an exact p value of 0.008, the smallest attainable with these group sizes. The same comparison against non-tumoural brain gave 0.55 in log2 units (p = 0.016), and adjustment for tumour size left 0.31 to 0.50 (p = 0.003 to 0.024). At the border the ratio of sphingomyelin to phosphatidylcholine rose by 0.79 in log2 units (p = 0.016, and 0.68 after adjustment for size, p = 0.013). The whole tumour and the border are two views of the same nine animals and the same pixels, at two spatial scales, and not two independent results. The measure reported for sphingomyelin in the whole tumour was chosen among four, of which the mean across species gives p = 0.11. Across the 17 class-level tests none survives Benjamini-Hochberg correction at 5 %, the lowest q value being 0.07.

## Discussion

Our results show that anti-PD-L1 given as a single agent reduces the volume of orthotopic GL261 tumours. At day 14, tumours of treated animals were 0.33 times the volume of control tumours in the IDH1-wild-type line (95 % CI 0.21 to 0.53; p < 0.001) and 0.42 times in the IDH1-mutant line (0.22 to 0.78; p = 0.006). The two lines were analysed separately, so each estimate stands on its own and nothing that follows compares them. Published work on checkpoint blockade in this model does not agree on what a single agent does. Anti-PD-1 alone gave no survival benefit in one orthotopic GL261 study [6]. In another study of the same model, single-agent anti-PD-1 and anti-PD-L1 produced long-term tumour-free survival in 50 % and 20 % of animals respectively [7]. Our endpoint is tumour volume followed over time rather than survival, so these results add a measurement to that disagreement rather than settling it.

Day 14 is a reference point of the model and not a day on which every animal was measured. In the IDH1-mutant line the confidence interval is wide, and these data fix the size of the effect there only loosely.

In the treated IDH1-wild-type tumours the apparent diffusion coefficient was higher, by 23 % over the whole tumour and by 29 % in the core (p = 0.027 and p = 0.030). The apparent diffusion coefficient rises as water moves more freely, which is what a tissue of lower cell density and a wider extracellular space would give, and it moved in the body of the tumour rather than at its border. Fractional anisotropy, which measures how much the movement of water prefers one direction over the others, fell by 24 % in the periphery of those same tumours (p = 0.005) and, in the IDH1-mutant line, by 25 % over the whole tumour and by 29 % in the core (p = 0.023 and p = 0.017). A fall in fractional anisotropy is a loss of directional organisation of the tissue. The two measures therefore report different things, one how freely water moves and the other how organised that movement is, and in these tumours they moved in different regions and in different lines.

Across the whole tumour, and relative to the healthy brain of the same section, the relative abundance of species assigned to sphingomyelin was higher in treated than in control animals, and phosphatidylcholine species shifted towards shorter chains (p = 0.016 for both; 0.008 and 0.004 after adjustment for tumour size). Sphingomyelin is made by transferring phosphocholine from phosphatidylcholine onto ceramide [16], so a shift in the balance between the two classes is the change that a move through that branch of membrane lipid metabolism would produce [17]. The ratios between classes, which do not depend on the normalisation, move in the same direction, and the same direction of change was present at the tumour border. The most economical reading is a remodelling of membrane lipid composition in the treated tumours. A change in the cell populations present in the tumour would give the same measurement, and that alternative is not hypothetical, since the brain tumour microenvironment carries an immune compartment of its own [18] and dendritic cells of tumour-bearing mice and of patients hold substantially more triglyceride than those of tumour-free hosts [19]. Free phosphocholine, choline and CDP-choline lie below the acquired mass range, so the part of choline metabolism on which the hypothesis of this project rests [14] is precisely the part these data cannot address.

Several limitations bound what this work can claim. Survival was not measured, and tumour volume is not a measure of benefit. The immune compartment was not characterised, no histology was performed, one antibody was tested as a single agent in one orthotopic model, and the lipid imaging comes from a single time point in nine animals of one line. The response did not differ between sexes in the IDH1-wild-type line (interaction p = 0.78), although the study was not sized for that comparison, and in the IDH1-mutant line the sex of the animals could not be separated from the order in which they were implanted. In patients, single-agent PD-1 blockade did not prolong survival over bevacizumab in recurrent GBM [2], and a volume measured in a mouse does not speak to that outcome.

## Figure legends

The fold changes in (C) are the values given in the text, expressed differently: 0.13, 0.54 and −0.40 in log2 units are +9 %, +45 % and −24 %. Two asterisks mark a class that differs before and after adjustment for tumour size, and the single asterisk on the long-to-short ratio marks one that differs only after that adjustment. Phosphocholine was not measured, because it falls below the acquired m/z range, so its box carries no value and the arrow leaving it is a hypothesis about the route rather than an observation. Ceramide carries no value either, because its summed class did not differ between the two groups (p = 0.25).

Absolute brightness is not comparable between sections, so the maps in (A) show the difference and the per-animal values in (B) and (C) measure it.

## References

1. Stupp R, Mason WP, van den Bent MJ, Weller M, Fisher B, Taphoorn MJB, et al. Radiotherapy plus concomitant and adjuvant temozolomide for glioblastoma. N Engl J Med. 2005;352(10):987–96. PMID: 15758009. doi:10.1056/NEJMoa043330

2. Reardon DA, Brandes AA, Omuro A, Mulholland P, Lim M, Wick A, et al. Effect of nivolumab vs bevacizumab in patients with recurrent glioblastoma: the CheckMate 143 phase 3 randomized clinical trial. JAMA Oncol. 2020;6(7):1003–10. PMID: 32437507. doi:10.1001/jamaoncol.2020.1024

3. Cloughesy TF, Mochizuki AY, Orpilla JR, Hugo W, Lee AH, Davidson TB, et al. Neoadjuvant anti-PD-1 immunotherapy promotes a survival benefit with intratumoral and systemic immune responses in recurrent glioblastoma. Nat Med. 2019;25(3):477–86. PMID: 30742122. doi:10.1038/s41591-018-0337-7

4. Berghoff AS, Kiesel B, Widhalm G, Rajky O, Ricken G, Wöhrer A, et al. Programmed death ligand 1 expression and tumor-infiltrating lymphocytes in glioblastoma. Neuro Oncol. 2015;17(8):1064–75. PMID: 25355681. doi:10.1093/neuonc/nou307

5. Nduom EK, Wei J, Yaghi NK, Huang N, Kong LY, Gabrusiewicz K, et al. PD-L1 expression and prognostic impact in glioblastoma. Neuro Oncol. 2016;18(2):195–205. PMID: 26323609. doi:10.1093/neuonc/nov172

6. Zeng J, See AP, Phallen J, Jackson CM, Belcaid Z, Ruzevick J, et al. Anti-PD-1 blockade and stereotactic radiation produce long-term survival in mice with intracranial gliomas. Int J Radiat Oncol Biol Phys. 2013;86(2):343–9. PMID: 23462419. doi:10.1016/j.ijrobp.2012.12.025

7. Reardon DA, Gokhale PC, Klein SR, Ligon KL, Rodig SJ, Ramkissoon SH, et al. Glioblastoma eradication following immune checkpoint blockade in an orthotopic, immunocompetent model. Cancer Immunol Res. 2016;4(2):124–35. PMID: 26546453. doi:10.1158/2326-6066.CIR-15-0151

8. Szatmári T, Lumniczky K, Désaknai S, Trajcevski S, Hídvégi EJ, Hamada H, et al. Detailed characterization of the mouse glioma 261 tumor model for experimental glioblastoma therapy. Cancer Sci. 2006;97(6):546–53. PMID: 16734735. doi:10.1111/j.1349-7006.2006.00208.x

9. Genoud V, Marinari E, Nikolaev SI, Castle JC, Bukur V, Dietrich PY, et al. Responsiveness to anti-PD-1 and anti-CTLA-4 immune checkpoint blockade in SB28 and GL261 mouse glioma models. Oncoimmunology. 2018;7(12):e1501137. PMID: 30524896. doi:10.1080/2162402X.2018.1501137

10. Louis DN, Perry A, Wesseling P, Brat DJ, Cree IA, Figarella-Branger D, et al. The 2021 WHO Classification of Tumors of the Central Nervous System: a summary. Neuro Oncol. 2021;23(8):1231–51. PMID: 34185076. doi:10.1093/neuonc/noab106

11. Kohanbash G, Carrera DA, Shrivastav S, Ahn BJ, Jahan N, Mazor T, et al. Isocitrate dehydrogenase mutations suppress STAT1 and CD8+ T cell accumulation in gliomas. J Clin Invest. 2017;127(4):1425–37. PMID: 28319047. doi:10.1172/JCI90644

12. Bunse L, Pusch S, Bunse T, Sahm F, Sanghvi K, Friedrich M, et al. Suppression of antitumor T cell immunity by the oncometabolite (R)-2-hydroxyglutarate. Nat Med. 2018;24(8):1192–203. PMID: 29988124. doi:10.1038/s41591-018-0095-6

13. Zhang L, Sorensen MD, Kristensen BW, Reifenberger G, McIntyre TM, Lin F. D-2-hydroxyglutarate is an intercellular mediator in IDH-mutant gliomas inhibiting complement and T cells. Clin Cancer Res. 2018;24(21):5381–91. PMID: 30006485. doi:10.1158/1078-0432.CCR-17-3855

14. Glunde K, Bhujwalla ZM, Ronen SM. Choline metabolism in malignant transformation. Nat Rev Cancer. 2011;11(12):835–48. PMID: 22089420. doi:10.1038/nrc3162

15. Wang HJ, Huang CY, Wei KC, Hung KC. A mass spectrometry imaging and lipidomic investigation reveals aberrant lipid metabolism in the orthotopic mouse glioma. J Lipid Res. 2022;63(12):100304. PMID: 36273646. doi:10.1016/j.jlr.2022.100304

16. Huitema K, van den Dikkenberg J, Brouwers JF, Holthuis JC. Identification of a family of animal sphingomyelin synthases. EMBO J. 2004;23(1):33–44. PMID: 14685263. doi:10.1038/sj.emboj.7600034

17. Hannun YA, Obeid LM. Sphingolipids and their metabolism in physiology and disease. Nat Rev Mol Cell Biol. 2018;19(3):175–91. PMID: 29165427. doi:10.1038/nrm.2017.107

18. Quail DF, Joyce JA. The microenvironmental landscape of brain tumors. Cancer Cell. 2017;31(3):326–41. PMID: 28292436. doi:10.1016/j.ccell.2017.02.009

19. Herber DL, Cao W, Nefedova Y, Novitskiy SV, Nagaraj S, Tyurin VA, et al. Lipid accumulation and dendritic cell dysfunction in cancer. Nat Med. 2010;16(8):880–6. PMID: 20622859. doi:10.1038/nm.2172

